# Clean lighting leads to decreased indoor air pollution and improved respiratory health in rural Uganda

**DOI:** 10.1101/455097

**Authors:** Viola N. Nyakato, Nicholas Mwine, Erez Lieberman Aiden, Aviva P. Aiden

## Abstract

Exposure to smoke is a major cause of respiratory illness in the developing world. To date, cookstoves have been the most widely studied source of smoke exposure in developing countries. We hypothesized that exposure to kerosene lighting, utilized by 86% of rural off-the-grid communities in sub-Saharan Africa may also be a significant source of smoke exposure and may be responsible for respiratory pathology. We performed an interventional field trial including 230 people in rural Uganda to assess the impact of clean lighting on indoor air pollution and respiratory health. Each member of the study households were asked about their exposure to smoke, the types of lighting they used, and their recent history of respiratory symptoms. Next, we provided solar-powered lamps to households in the intervention group, and compared to households in the control group who continued to use kerosene lamps. We monitored indoor air quality in a subset of intervention and control households over a three-month period, and performed an exit survey to assess symptoms of respiratory illness in both groups. All of the households we surveyed were found to use kerosene lamps as their primary lighting source. We found that the average person was exposed to 3.3 hours of smoke from kerosene lamps, as compared to 44 minutes of exposure from cookstoves. Next, we found that average soot levels (elemental carbon) in intervention homes were 19-fold lower than soot levels in control homes. After three months, we observed reduced rates of all symptoms assessed, and significantly reduced risk of cough, sore throat, and overall illness in the intervention homes. Our findings demonstrate that kerosene lighting is a significant source of smoke exposure in the developing world, and that the introduction of clean lighting in homes reliant on kerosene lighting can have a rapid and significant impact on overall health.

## Introduction

For decades, respiratory infections have been the number one killer of children under the age of five worldwide.^(1)(2)(3)^ They are also a significant contributor to morbidity and mortality in other age groups.^(4)^

In developing nations, much of the air pollution – especially indoors – stems from the burning of fuels based on petroleum and biomass by individual households. Such fuels are used for cooking, heating, and lighting by over half the world population.^(5)^

Most studies exploring indoor air pollution in the developing world have focused on smoke exposure due to cooking.^(6)(7)^ The most extensive study to date was the RESPIRE trial, which examined the impact of cleaner-burning cookstoves on 518 young children in Guatemala. Mothers in the intervention group received a cookstove that produced less pollution than traditional stoves. The respiratory health of the intervention group showed improvements in rates of serious pneumonia at the end of the 18 month trial.^(7)^

We suspected that lighting, and specifically kerosene-based lamps, are also a significant cause of morbidity in the developing world. Cooking is usually done by adult women in a well-ventilated kitchen that is separate from the main part of the house. In contrast, indoor lighting typically: (i) affects all members of the household, (ii) is used in the more poorly-ventilated main house, and (iii) is produced using kerosene lamps, which generate a great deal of smoke.

We performed an interventional field trial in rural Uganda to assess the impact of clean lighting on respiratory health. Our field trial examined 230 people drawn from 50 households. At the outset, participants completed health and smoke exposure surveys. Households were randomized to intervention and control groups. The intervention households then received smoke-free solar lights and the control households did not (they received lights as a gift at the end of the study). During the three month intervention, we monitored fine particle (PM2.5) pollution in a subset of households. After three months, health surveys were again administered to all 230 people.

We showed that, prior to our intervention, the average subject reported nearly five times as many hours of smoke exposure due to indoor lighting as compared to cooking. During the trial, our analysis of airborne particulates revealed a 95% decrease in average fraction of elemental carbon (p value <0.0001) In exit surveys, the intervention group reported lower rates of all five symptoms of respiratory illness studied, with a statistically significant reduced risk in three: cough, sore throat, and general illness.

## Methods

## Household Selection

We obtained IRB approval for our study from Harvard University, the Mbarara University of Science and Technology (MUST), and the Ugandan National Council for Science and Technology (UNCST). We also described the study design to the local community at town hall meetings, and obtained permission to proceed from the community’s leadership.

Fifty study households were selected by lottery from attendees at the town hall meetings of two parishes in the Mbarara region of Uganda. Households where members reported access to electricity via either the electrical grid or solar panels were to be excluded from the study; in fact, no household among the fifty chosen reported such access. Half of the households were assigned to the intervention group, and half to the control group. Households in the intervention group received a solar light (specifically, a d.light S10 solar lantern), and were asked to use these instead of their kerosene lamps during the study period. Kerosene lamps were not removed from the intervention households. Households in the control group were asked to continue utilizing their normal sources of indoor lighting, and were informed that they would receive a solar light at the end of the study.

Typical homes were 3 room enclosures, comprising a living room – the primary room in which lighting was used – as well as a bedroom and a storeroom (Figure 1A). A separate building was used as a kitchen. Cooking exposure was limited to the kitchen building, which tended to be less enclosed (e.g. no door, unpaned windows – Figure 1B).

**Figure 1.**
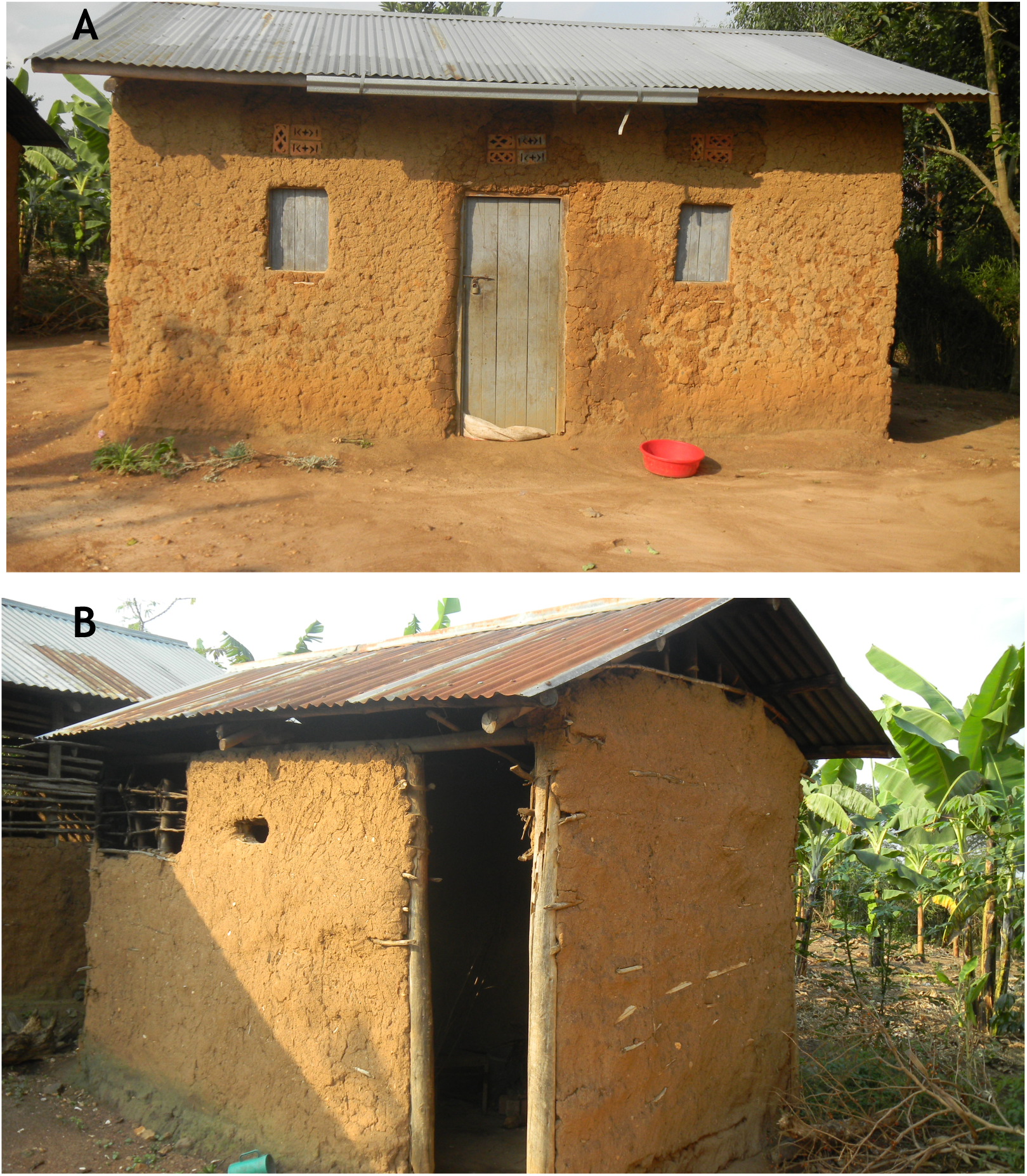
Typical home in the study communities in rural Uganda. **A)** Homes typically contained a living room where lighting was used, a bedroom, and a storeroom. Doors and windows were usually closed, offering limited ventilation when using kerosene lighting. **B)** Households typically maintained an outbuilding for cooking. These buildings were more open than the homes themselves, and generally did not have doors or window covers that closed.

A field worker visited each home every 7-10 days.

## Air Quality Measurements

We evaluated particulate matter in eight homes, four from the intervention group and four from the control group. We assessed levels of 2.5 µm particles using Pall Corp. 25 mm quartz filters mounted in URG filter samplers with 2.5 µm cyclones. Four filter samplers were rotated between the eight homes. The filter samplers were placed approximately 1.7 m from the ground in each dwelling, and the filters were collected approximately every 3 days. Filters were returned to the US and analyzed for total, organic and elemental carbon (“TC,” “OC” and “EC”, respectively) using a Sunset Labs OCEC Analyzer and the NIOSH 870 protocol. On average, each home assessed was monitored for 12 days over the course of the study. To assess ambient levels of particulate matter, filter samplers were placed at a similar height outdoors in the parish.

## Symptom Assessment

At the outset of the study, each member of the 50 selected households was asked to complete a survey. The survey asked for demographic information, estimates of the number of hours spent cooking and using artificial lighting in a typical week, a description of the fuel sources used for both activities, and symptoms of illness during the past three months. The survey was originally composed in English, translated into Runyankole by a student at MUST, and administered verbally, in person, by a native speaker of Runyankole. The survey is available in both languages as part of the supplemental online materials. For those too young to complete the survey themselves, an adult member of their household completed the survey on their behalf. At the conclusion of the study, a similar survey was administered. Only the responses of people who had completed both the entry and exit surveys were included in our analysis.

There are several potential sources of bias in our survey methodology. Because participants were informed about the purpose of the study, a placebo effect could be present. There could also be a bias towards greater consciousness of symptoms due to the Hawthorne effect: participants may take particular notice of symptoms because they know that they will be asked to report these symptoms at the end of the study. Additionally, because the entry and exit survey questions refer to a period of time spanning roughly six months, seasonal effects may have contributed to the differences observed. However, given the spatial proximity of the study households, we believe that any such effect would have had a similar impact on both the control and intervention groups. The retrospective study design described above was chosen in order to reduce the amount of bias.

All data was analyzed by researchers who were blinded as to whether the respondent was in the control or intervention group.

## Results

In total, 230 people in 50 households completed both the entry and exit surveys. 123 people were in the intervention group, representing all 25 households. 107 people were in the control group, again representing all 25 households.

Notably, the entry survey revealed that most of the study population reported no exposure to cook-smoke, but several hours of smoke exposure per day due to indoor lighting (Figure 2). The mean and median cook-smoke exposure was 44 minutes and 0 minutes, respectively; the mean and median values for lighting smoke were 3.3 hours and 4 hours. Each of the 50 households reported utilizing kerosene for most or all of their lighting needs. At least one member of twenty different households reported using a battery flash-light or candles as a supplemental light source. In every case, the alternate forms of lighting were reported to be used “some/a little bit,” and kerosene is listed as the primary source of lighting fuel. Every household used firewood as its primary source of cooking fuel, with members of 5% of households using “some/a little bit” of charcoal as well. 9·3% of adults (individuals over 18 years old) reported that they were smokers. This value is consistent with other studies done in the region.^(8)^

**Figure 2.**
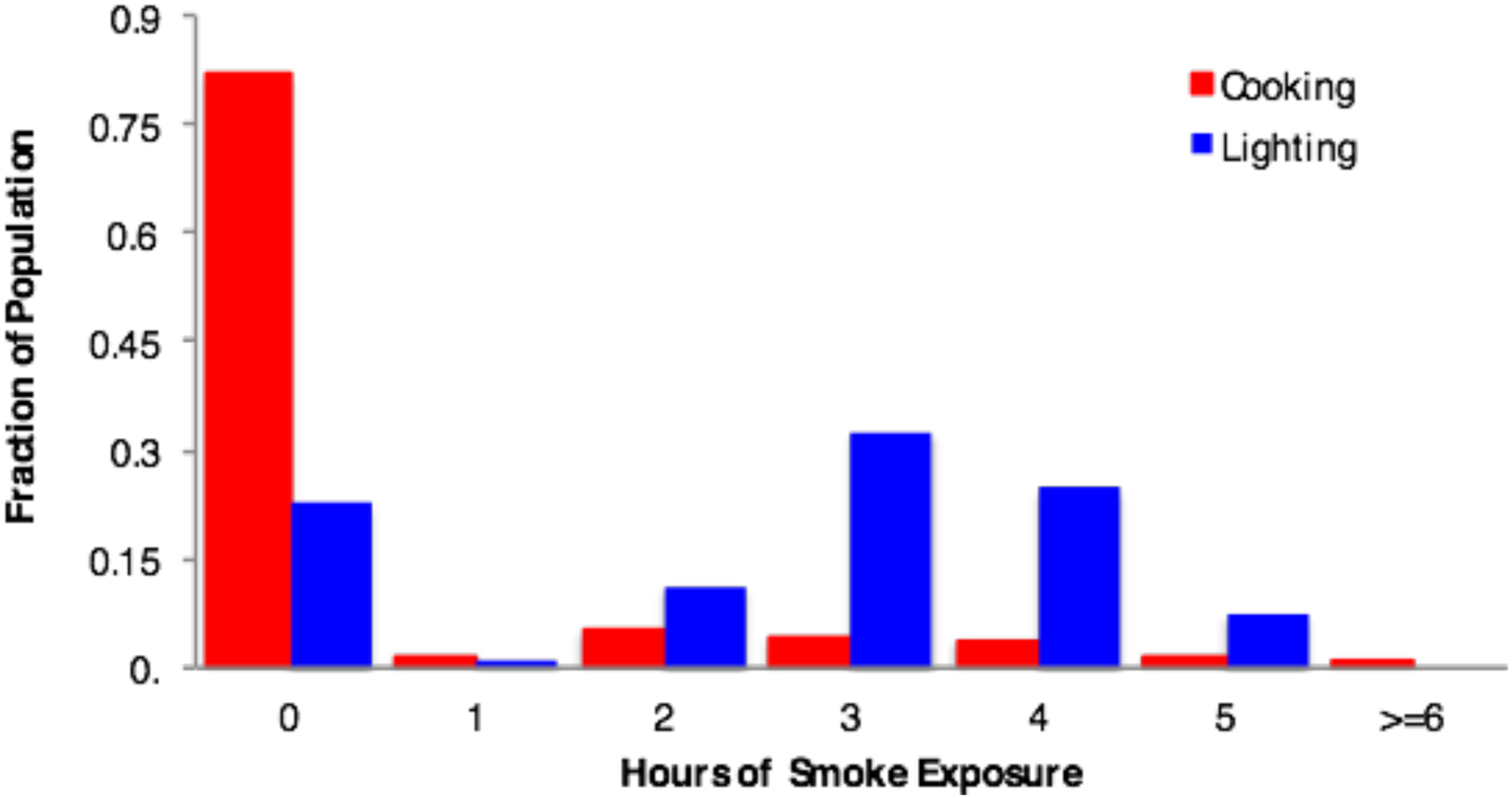
Average per person exposure time to smoky lighting is nearly 5-fold higher than average per person exposure time to cooking smoke. ~80% of household members are not exposed to cooking smoke at all, but > 70% of household members are exposed to at least 2 hours of kerosene lighting daily. Every household in the study listed burning kerosene as the primary source of lighting in the home.

## Air Quality

Strikingly, elemental carbon was reduced by 95% on average in intervention homes as compared to control homes. The difference in soot levels was immediately apparent upon visual examination of the filter (Figure 3A). Fractions of elemental carbon (soot) in intervention homes were five-fold lower than in control homes (P-value <0·0001 using t-test). As expected, there was no significant difference between levels of airborne organic carbon in control and intervention homes (Figure 3B).The mean total carbon was reduced from 20 µg/m3 in the control homes to 7 µg/m3 in the intervention homes.

**Figure 3.**
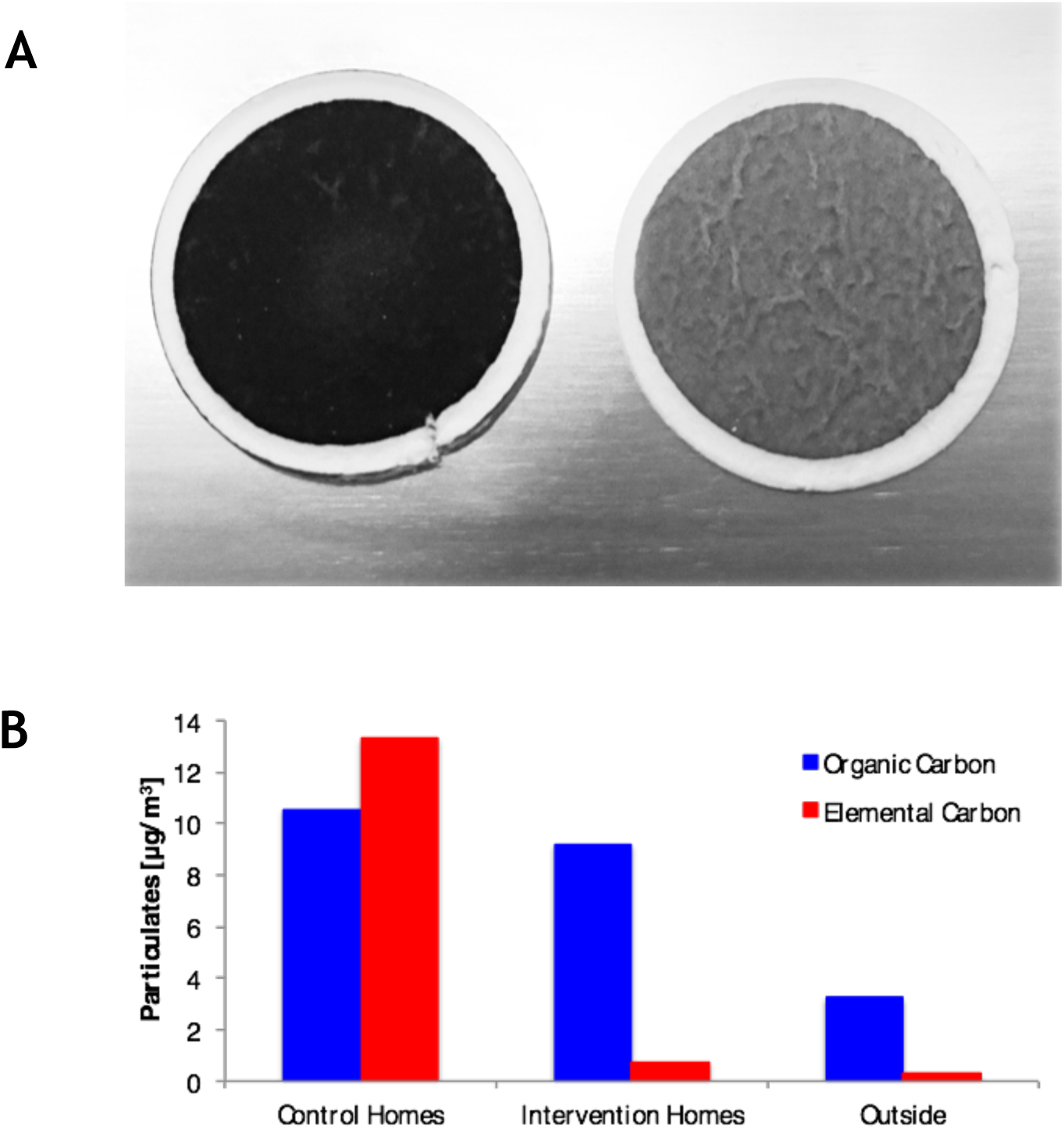
Clean lighting reduces indoor particulate air pollution. **A)** On visual inspection, filters capturing 2.5 µm particles over a 3 day period show markedly more particulate material in the control homes as compared to the intervention homes. **B)** Evaluation of all filters from control vs intervention homes shows dramatic reduction in elemental carbon (soot).

## Symptomatology

At the beginning of the study, there were no statistically significant differences in the symptoms reported by the control and intervention groups (Table 1).

**Table 1.**

|  | **Intervention Group** | **Control Group** | **P-value (Fisher Exact Test)** |
| --- | --- | --- | --- |
| **Total Number of People** | 123 | 107 |  |
| **Sick** | 87 | 84 | 0.226 |
| **Cough** | 69 | 49 | 0.146 |
| **Difficulty Breathing** | 28 | 30 | 0.366 |
| **Wheezing** | 22 | 25 | 0.329 |
| **Sore Throat** | 25 | 27 | 0.430 |

At the end of the study, rates of all five symptoms evaluated were lower in the intervention group (Table 2). Both the reduction in rate of general illness and the reduction in rate of sore throat were statistically significant after Bonferroni correction. The reduction in rate of cough yielded a P-value of 0.026, but with confidence intervals 0.57-0.90. (Figure 4).

**Table 2.**

|  | **Intervention Group** | **Control Group** | **P-value (Fisher Exact Test)** |
| --- | --- | --- | --- |
| **Total Number of People** | 123 | 107 |  |
| **Sick** | 62 | 72 | 0.0069 |
| **Cough** | 41 | 50 | 0.026 |
| **Difficulty Breathing** | 23 | 22 | 0.42 |
| **Wheezing** | 16 | 17 | 0.33 |
| **Sore Throat** | 13 | 25 | 0.0075 |

**Figure 4.**
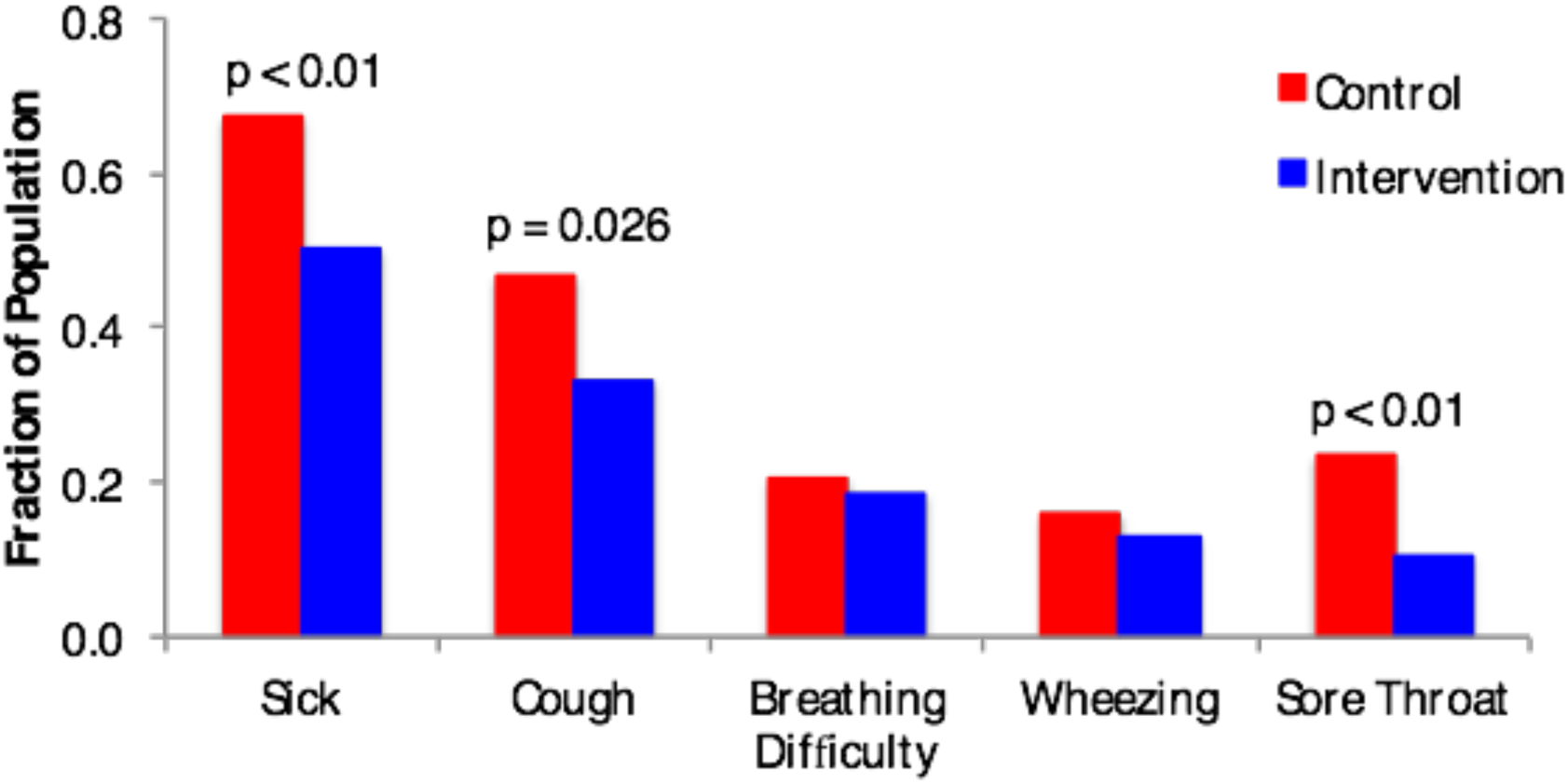
Clean lighting improves respiratory symptoms. Introducing clean lighting for 3 months significantly reduced the incidences of illness, cough, and sore throat as compared to the incidences of those symptoms in control homes over the same period.

Due to the sample size, our ability to obtain meaningful results when examining sub-groups was limited. However, we note that the reduction in the overall rate of illness in children (individuals younger than 18) showed a statistically significant decline (P<0.05).

## Discussion & Conclusions

Respiratory diseases lead to nearly two million preventable deaths each year in developing nations, where they are the number one cause of death in children under five. ^(1)(9)(10)^ Lower respiratory infections account for roughly 11% of all deaths in low-income countries, and 4-5% of deaths in middle and upper income countries as well.^(4)^

Because of the landmark work of Smith *et al.,* ^(7)^ the causal role of stove smoke in health – the so-called “killer in the kitchen” – is now widely recognized. In contrast, kerosene lamps are the primary source of light for over 1.7 billion people worldwide,^(11)^ but the health impact of the smoke they produce has not been explored.

Our study was performed in rural Uganda, a country whose per capita GNI of $1,310 falls at the very low end of the middle-income range. Based on our findings, smoke exposure due to kerosene-based lighting far exceeds smoke exposure due to any other household source; in particular, the typical individual is exposed to nearly five-fold more hours of smoke from lighting than from cooking. The results of our trial suggest that replacing kerosene lanterns with clean lighting leads to reductions in overall rates of illness, cough, and sore throat among all members of the household – not just those exposed to cook smoke. For instance, by the end of our three-month trial, we observed a 40% reduction in reported cough symptoms in the intervention group, as compared to a 2% increase in the control group. Our work underscores the potential positive impact of clean lighting on objective air pollution measures as well as on symptoms reported by members of the household.

Our study did not assess the relative impact of the various types of smoke exposure per unit time; it is possible, for instance, that an hour of exposure to cook smoke causes more harm than an hour of exposure to smoke from a kerosene lamp. (The reverse is also possible.) We also did not quantify the relationship between duration of exposure and health outcomes. Both of these questions warrant additional study. Future studies would benefit from a larger group size, monitoring of additional pollutants and symptoms, objective (rather than subjective) clinical assessments, and a longer trial. In particular, a multi-year trial would better account for seasonal variation, as well as inter-year variation in severity of annual illnesses, such as influenza, or respiratory syncytial virus.^(12)^

Despite the need for additional research, the findings of the present study are highly suggestive. Taken together, they show that the health burden of kerosene lamps is very significant. From the technological standpoint, lighting homes cheaply and cleanly is a solved problem. Numerous technologies have been developed whose stated purpose is to provide better lighting options to people with limited financial resources and no access to an electrical grid. (The d.lights we employed in this study are one such example.) The challenge that remains is to make these devices more widely available.^(13)^ Our study suggests that improving the availability of clean lighting could improve health for nearly two billion people.

## Acknowledgements

We would like to acknowledge the contributions of Judith Iradukunda (MUST), Samuel Killewo (Harvard), Wendo Mlahagwa (MUST), and Jonathan E. Rea (MIT) for their contributions to the field study data collection and Kelli Fischer (UCB), who processed the filters.

Authors report no personal or financial conflicts of interest in this study

VNN assisted in the design of the clinical component and led the field implementation

NW assisted in the implementation of the field studies

NM assisted in the design, implementation and analysis of the air quality component

ELA assisted with the design of the field study and writing the paper.

APA conceived the study, designed the clinical and air quality components of the field study, analyzed the field and air quality data, and wrote the paper.

---

**Supplemental Tables 1A-C:**
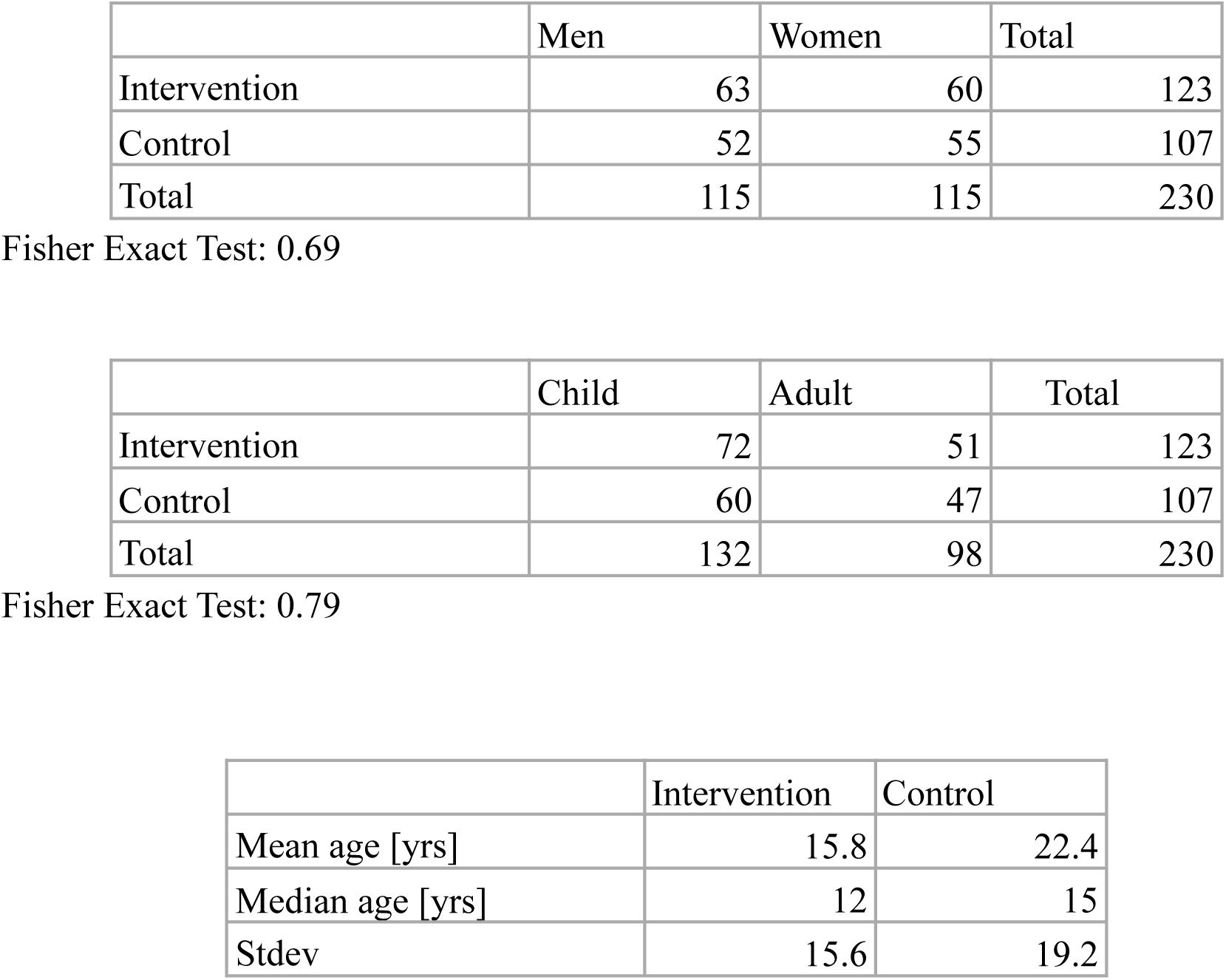
Comparison of demographics in control vs intervention group. In terms of distribution of men and women, and children and adults, the groups were comparable. When evaluating the mean/median ages, it is important to note that in 17 households, the male head of household declined to state his age. These men were listed as adults. These households were predominantly in the Intervention group. When analyzing the age distribution, the median age of the intervention group appeared lower due to the absence of ages of a significant number of adults.

